# Multiomics Analysis Reveals Extensive Remodeling of the Extracellular Matrix and Cellular Metabolism Due to Plakophilin-2 Knockdown in Guinea Pigs

**DOI:** 10.1101/2024.03.11.584401

**Authors:** Rui Song, Haiyan Wu, Lihui Yu, Jingning Yu, WenHui Yang, WenJun Wu, Fei Sun, Haizhen Wang

**Affiliations:** College of Veterinary Medicine, Yunnan Agricultural University, #452 Fengyuan Road, Kunming 650021, China; College of Tea Science, Yunnan Agricultural University, #452 Fengyuan Road, Kunming 650021, China; Department of Ultrasoned, the Second Affiliated Hospital of Kunming Medical University, #376 Dianmian Avenue, Wuhua District, Kunming 650032, China; Department of Cardiology, the Affiliated Hospital of Kunming University of Science and Technology, the First People’s Hospital of Yunnan Province, #157 Jinbi Road, Kunming 650032, China; Department of Cardiology, Fuwai Yunnan Cardiovascular Hospital, #528 Shahe North Road, Kunming 650102, China

**Keywords:** Arrhythmogenic right ventricular cardiomyopathy, sudden cardiac death, extracellular matrix, guinea pigs

## Abstract

Arrhythmogenic right ventricular cardiomyopathy (ARVC) is a leading cause of sudden cardiac death (SCD) in young individuals, yet the mechanisms underlying its pathogenesis, particularly the role of plakophilin-2 (PKP2), remain incompletely understood. This study aimed to elucidate the profile of molecular and metabolic consequences of PKP2 knockdown in a guinea pig model of ARVC. We employed adeno-associated virus serotype 9 (AAV9) to deliver PKP2 shRNA, establishing a model that recapitulates key features of human ARVC, including right ventricular (RV) enlargement, sudden death, and cardiac lipid accumulation. Proteomic analysis revealed significant dysregulation of extracellular matrix (ECM) proteins, PI3K-Akt signaling, and energy metabolism in PKP2-deficient RVs. Metabolomic profiling further highlighted alterations in lipid metabolism and inter-metabolites of TCA cycle, with a notable shift towards fatty acid oxidation. These findings suggest that PKP2 deficiency triggers a cascade of molecular events leading to ECM remodeling, metabolic reconfiguration, and potential mitochondrial dysfunction, which may contribute to the development of ARVC. Our study provides novel insights into the early molecular mechanisms of ARVC and identifies potential therapeutic targets for this underexplored disease.

## Introduction

ARVC is a significant cause of SCD, accounting for up to 20% of SCD cases in young population^1,2^. Additionally, ARVC Patients without SCD often develop dual ventricle failure^3,4^. The prevalence of ARVC ranges from 1 in 5000 to 1 in 3000 depend on the population studied. 60% of ARVC have identifiable genetic causes, with mutations in desmosomal proteins being the most common, particularly in PKP2^5,6^.

Diagnosis of ARVC still challenged and typically relies on a combination of structural and functional examinations, familial characteristics, echocardiography, and electrocardiography (ECG) findings^7,8^. However, the early stages of ARVC are often asymptomatic, and SCD can be the initial manifestation^9^. Therefore, it is likely that the frequency of ARVC is underestimated. Consequently, gaining a comprehensive understanding of mechanical details of early ARVC is crucial for prevention of ARVC caused SCD and heart failure.

Typical ARVC related pathological changes include fibrofatty replacement and RV dilation. Many ARVC patients remain asymptomatic during the first decade of life, with symptoms typically emerging in the second or third decade^2^. This latency suggests that early metabolic remodeling may play a pivotal role in the progression of ARVC. Nevertheless, the specific molecular pathways and metabolic alterations that arise from desmosomal gene mutations, particularly following PKP2 knockdown, and their contributions to ARVC pathogenesis, are not fully understood.

To address this question, a novel ARVC larger animal model is required. While mouse models have been instrumental in ARVC research, they present certain limitations. Firstly, mice are relatively small, which poses challenges for echocardiography without specialized and expensive equipment. Secondly, mice may not fully develop the classic symptoms of cardiomyopathy observed in humans, as evidenced by some studies^10^. Therefore, it is imperative to develop a large animal model that not only accommodates standard echocardiography and ECG equipment but also mirrors the genetic defects associated with human ARVC.

The guinea pig has been recognized as a suitable model for cardiovascular diseases^11,12^. Its size allows for the evaluation of cardiac function and structure using standard echocardiography and ECG machines^13^. Therefore, we developed an ARVC guinea pig model by employing an adeno-associated virus serotype 9 (AAV9) vector to deliver small hairpin RNAs (shRNA) targeting plakophilin-2 gene (*pkp2*). Subsequently, we conducted proteomic and metabolomic analysis to assess protein expression and plasma metabolites profiles.

## Results

### Establishment of ARVC guinea pig model

To establish a robust animal model for investigating ARVC, Guinea pigs were selected as the experimental model and injected with AAV9 carrying PKP2 shRNA (PKP2i groups) via the forelimb veins, while the control group received AAV9 carrying non-targeting shRNA (CON groups). Echocardiography was performed monthly to evaluate the cardiac structure and function of the guinea pigs (Figure 1A). Each group consisted of 8 male and 8 female guinea pigs. After a 4-month period following AAV9 injection, the guinea pigs were euthanized, and samples were collected from the RV. The knockdown efficiency of PKP2 was determined through western blot experiments, which revealed a significant decrease in PKP2 protein levels in the knockdown samples (Figure 1B). Notably, there were no significant differences in body weight or the ratio of heart weight to body weight between the control and PKP2i groups (Figure 1C, D). Strikingly, similar to observations in human patients, sudden death cases were observed in guinea pigs with reduced PKP2 expression, with 2 males and 1 female guinea pig experiencing sudden death accompanied by right ventricular enlargement (Figure 1E). This finding coincides with clinical observations in humans and underscores the relevance of our animal model in studying ARVC.

**Figure 1.**
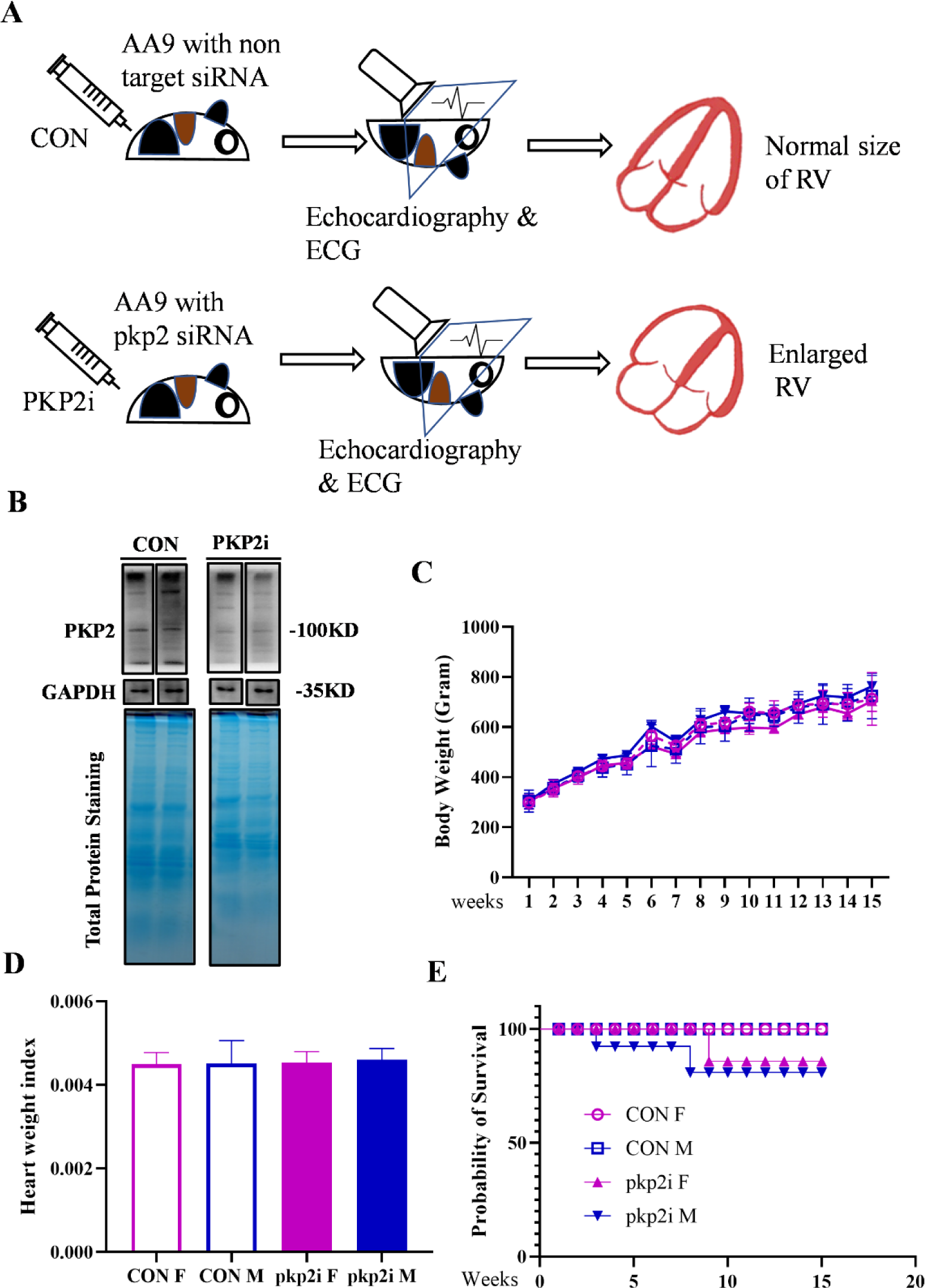
Construction of an ARVC guinea pig model. (A) Schematic of the experimental procedure used to establish an ARVC guinea pig model utilizing AAV9 vectors. (B) Western blot analysis of PKP2 and Coomassie staining for total protein in right ventricular samples. CON represents guinea pigs injected with AAV9 expressing non-targeting shRNA (n=16, 8 males and 8 females), while PKP2i represents guinea pigs injected with AAV9 expressing PKP2 shRNA (n=16, 8 males and 8 females). (C) Body weight curves over time for CON and PKP2i groups. (D) Ratio of heart weight to body weight for CON and PKP2i groups.

### Structural and functional alteration of right ventricles when PKP2 knocks down

To assess the impact of PKP2 knockdown on the structure and function of the RV in guinea pigs, we performed monthly echocardiography following AAV9 injection. Imaging analysis revealed a significant enlargement of the right ventricle in male guinea pigs of the PKP2i group starting from the second month post-injection, compared to the CON group. Notably, structural alterations in the RV of female guinea pigs were observed from the third month post-injection, compared to the CON group (Figure 2A).

**Figure 2.**
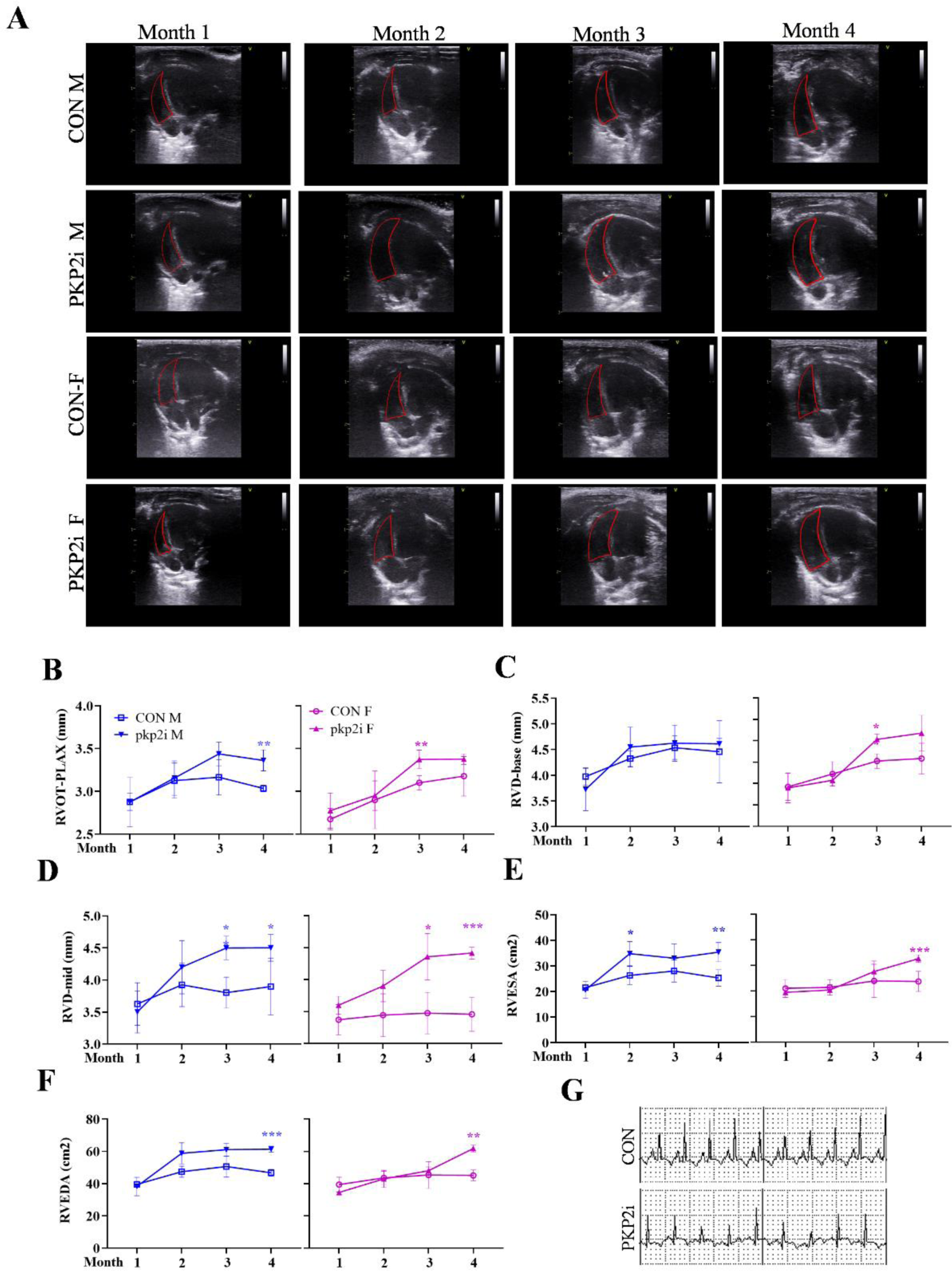
Echocardiographic evaluation of right ventricular structure. (A) Representative echocardiographic images of male (M) and female (F) guinea pigs from 1 to 4 months post-AAV9 injection. Quantitative analysis of (B) RVOT proximal long axis (RVOT-PLAX), (C) RV basal diameter (RVD-base), (D) RV median diameter (RVD-mid), (E) RV end-systolic area (RVESA), and (F) RV end-diastolic area (RVEDA) over time. (G) ECG curve of CON and PKP2i guinea pigs. Data are mean ± SD, n=8/group. Statistical comparisons were made by unpaired Student’s t-test; P<0.05, *P<0.01, **P<0.001.

One month after AAV9 injection, no significant differences were observed in RV structural parameters. However, at two months post-injection, the male PKP2i group exhibited a significant increase in RVD-mid, RVEDA, and RVESA indices compared to the male CON group. In contrast, no significant differences were observed in the female PKP2i group. By the third month post-injection, RVOT-PLAX showed a significant increase in the male PKP2i group compared to the CON group. Additionally, the female PKP2i group exhibited significant increases in RVOT-PLAX, RVD-basal, RVD-mid, RVEDA, and RVESA compared to female controls (Figure 2B-F). However, no significant differences were observed in left ventricular functional parameters between the PKP2i and CON groups (Figure S1). ECG examination also showed normal results at four months post-injection (Figure 2G).

### Enlarged RV and abnormal mitochondrial structure

After a 4-month period following AAV9 injection, we observed a significant enlargement of RV in both male and female guinea pigs in the PKP2i group. Subsequently, the guinea pigs were sacrificed, and the anatomical examination of the heart confirmed the enlargement of the RV space, consistent with the echocardiography findings (Figure 3A). Histological analysis of heart tissue sections from the PKP2i group revealed thinner RV walls. Sirius red staining demonstrated that the level of fibrosis was comparable to the CON group in the PKP2i group (Figure 3B). Though the triglyceride concentration was mild but significantly increased in the RV samples from PKP2i group (Figure 3C).

**Figure 3.**
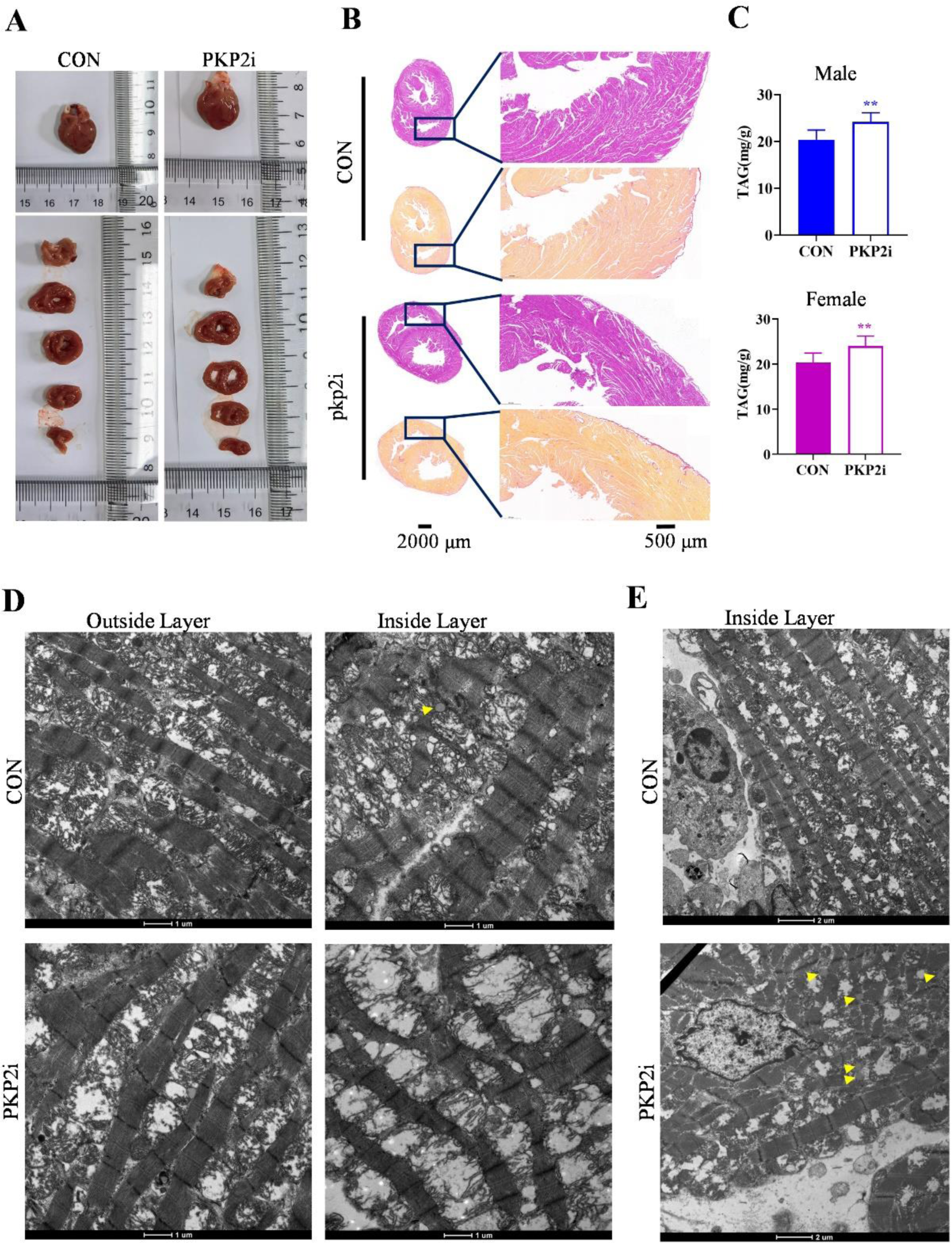
Structural alterations in the right ventricle of PKP2-deficient guinea pigs. (A) Gross morphology of right ventricle (RV) from representative CON and PKP2i guinea pigs. (B) Hematoxylin & eosin (H&E) and Sirius red staining of RV sections. Scale bars = 2000 μm (left), 500 μm (right). (C) Triglyceride content (TAG) in RV samples from male and female guinea pigs. (D-E) Transmission electron microscopy (TEM) images of the outer and inner layers of the RV wall from CON and PKP2i groups. Scale bars = 1 μm (left), 2 μm (right). Arrowheads indicate lipid droplets.

To investigate the impact of PKP2 knockdown on the two layers of RV muscle, we isolated and sampled the outer and inner layers of the guinea pig RV muscle. These samples were then subjected to transmission electron microscopy (TEM) analysis. The results indicated morphological changes in the mitochondria of both layers of the RV muscle in guinea pigs with reduced PKP2 expression. Compared to the CON group, the PKP2i group exhibited a higher prevalence of swollen mitochondria (Figure 3D). Among the male guinea pigs in the PKP2i group, two individuals displayed severe mitochondrial swelling in the inner layer of the RV muscle. Furthermore, other PKP2i guinea pigs with mild mitochondrial swelling exhibited an increased content of lipid droplets (Figure 3E). These findings were consistent with the biochemical examination of the RV muscle wall, which revealed a 20% increase in triglyceride content in the PKP2i group (Figure 3C). Thus, we successfully established the guinea pig model of arrhythmogenic right ventricular cardiomyopathy (ARVC). The focus now turns to understanding the underlying mechanisms within the RV muscle wall of the PKP2i group.

### Differential Proteomics and Functional Pathway Analysis in PKP2 Knockdown RV Muscle Wall

To unravel the intricate molecular changes occurring in the RV muscle wall caused by *pkp2* knock down, we conducted a comprehensive DIA (Data independent acquisition) quantitative proteomics study. Utilizing principal component analysis (PCA), we observed distinct distribution patterns of samples from the CON and PKP2i groups, indicating proteomic alterations in the PKP2i samples compared to the CON group (Figure 4A). Subsequently, we conducted an in-depth analysis of the proteomics data, detecting a total of 7800 proteins. Using Gene Ontology (GO) analysis, we determined the membrane proteins accounted for 43.38%, then the second abundant part was Cytoplasm proteins, accounted for 29.1% (Figure 4B). Next, 268 differential expressed proteins were identified. Among the significantly differential proteins, 170 of them were upregulated and 98 of them were downregulated (Figure 4C). The heat map showed the profile of differential proteins (Figure 4D). According GO analysis, 47.54% of differential proteins were identified as membrane proteins, 22.95% and 19.67% were cytoplasm and extracellular region proteins respectively (Figure 4E).

**Figure 4.**
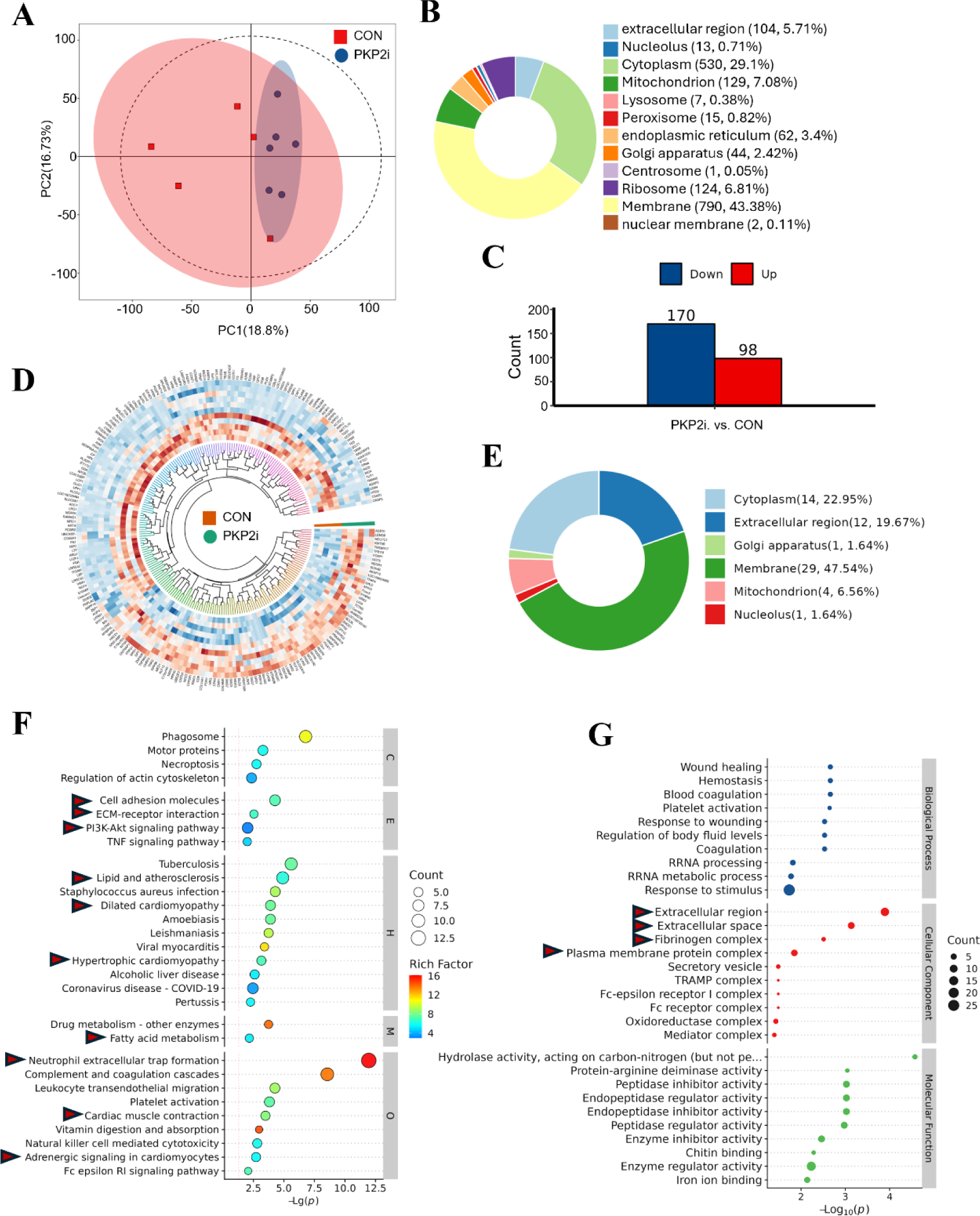
Differential protein expression profiling of PKP2i and CON right ventricle samples. (A) Principal component analysis (PCA) of CON and PKP2i proteomic datasets. (B) Graphical representation of the subcellular distribution of detected proteins, categorized by their cellular location. (C) Bar chart depicting the counts of upregulated (red) and downregulated (blue) proteins identified as significantly differentially expressed between PKP2i and CON samples. (D) Circular heatmap displaying the expression patterns of differentially expressed proteins, with color intensity corresponding to the level of expression. (E) Subcellular distribution of significant differential proteins, highlighting the changes in protein localization in PKP2i compared to CON samples. (F) Kyoto Encyclopedia of Genes and Genomes (KEGG) and (G) Gene Ontology (GO) pathway analysis, with red arrows indicating pathways of particular interest that are affected by PKP2 knockdown. Proteomics data were analyzed using Perseus, Excel, and R software. Hierarchical clustering used Euclidean distance and the complete agglomeration method with the heatmap R package. Enriched pathways were considered significant at p<0.05.

In order to elucidate the potential impact of the 268 differential proteins, we conducted Kyoto Encyclopedia of Genes and Genomes (KEGG) and Gene Ontology (GO) analyses encompassing all these proteins. Our KEGG analysis revealed a significant alteration of pathways associated with extracellular matrix (ECM), including cell adhesion molecules, ECM-receptor interaction and neutrophil extracellular trap formation (Figure 4F). Some important cardiac metabolic pathways altered following the knock down of *pkp2* as well, including PI3K-Akt pathway, lipid and atherosclerosis, fatty acid metabolism pathways. Dilated, hypertrophic cardiomyopathy associated pathways, adrenergic signaling in cardiomyocytes and cardiac muscle contraction associated pathways altered as well (Figure 4F).

Coincident with KEGG analysis, GO analysis revealed alteration of multiple plasma membrane complex pathways, including extracellular region, extracellular space, fibrinogen complex and plasma membrane protein complex (Figure 4G).

### Extracellular Matrix (ECM) Remodeling Post-PKP2 Knockdown: Implications for ARVC

The ECM is a critical determinant of cardiac structure and function. In this study, we have elucidated the molecular mechanisms underlying ECM remodeling following the knockdown of PKP2. Our analysis, informed by GO and KEGG, have identified the ECM as a central hub in the molecular pathogenesis of ARVC.

We observed a significant upregulation of Foxo3, a key transcription factor in ECM degradation, in the ARVC cohort (Figure 5A). This was contrasted by the downregulation of liver-derived ECM components, including fibronectin (FN1) and fibrinogen alpha and beta chains (H0V0Z9 and FGB), in the PKP2i group (Figure 5B-D). Additionally, bone marrow-derived ECM components such as extracellular matrix protein 1 (ECM1) and dermatopontin (DPT) showed reduced expression following PKP2 knockdown (Figure 5E, F).

**Figure 5.**
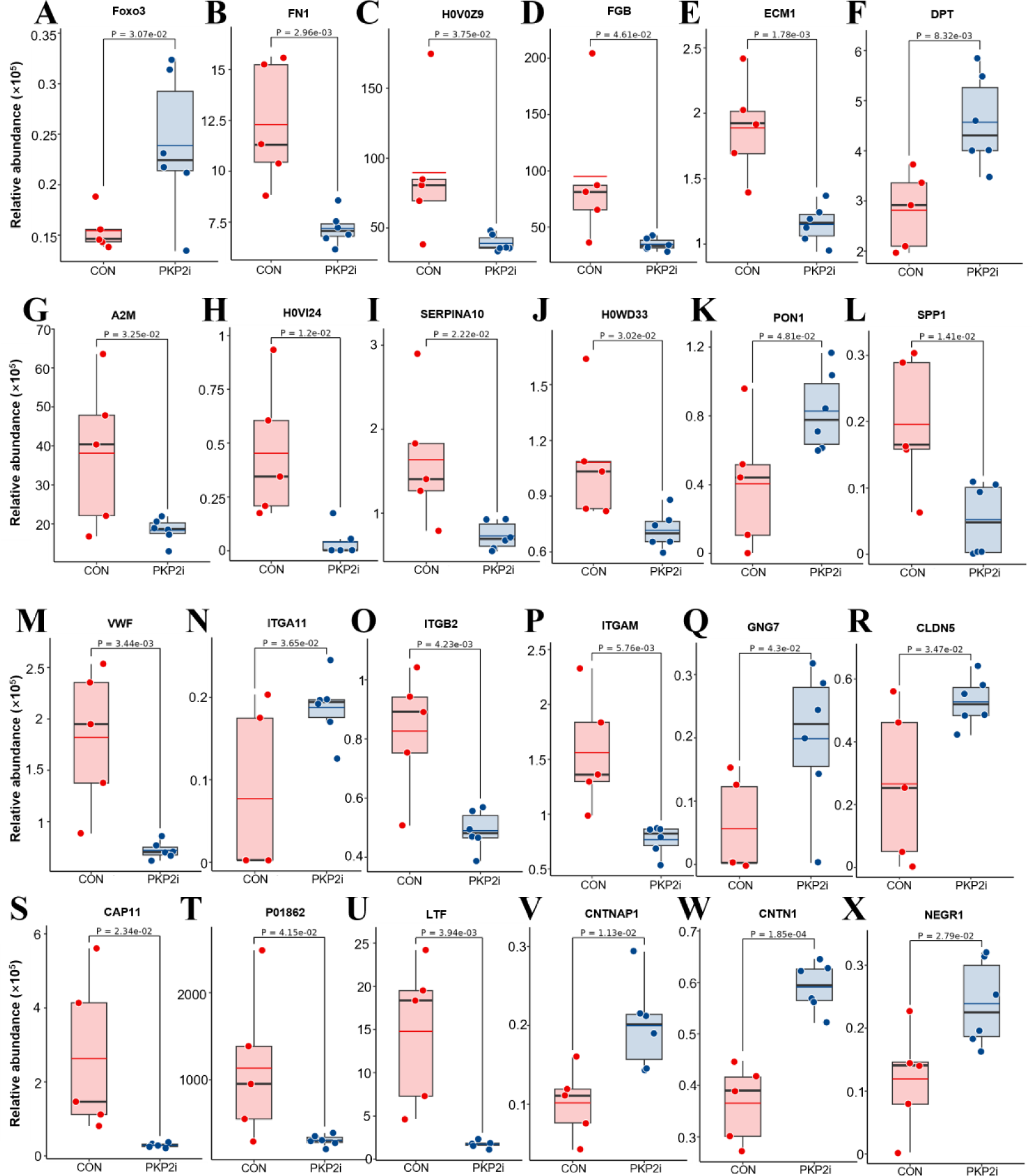
Altered ECM Protein Expression in PKP2-Deficient Right Ventricles. The differential expression analysis of ECM proteins in right ventricle samples from CON and ARVC animals, which are representative of PKP2 deficiency. Data are expressed as mean relative abundance ± standard deviation (SD), with n=5 for the CON group and n=6 for the PKP2i group. The Y-axis denotes the relative abundance of proteins, scaled by a factor of 10^5^. Statistical significance, as determined by P values, is indicated for each protein graph.

ECM components with protease inhibitor activity, including alpha-macroglobulin (A2M) and serpin family members (H0VI24, SERPINA10, H0VWD33), were notably decreased in the PKP2i group (Figure 5G-J). This downregulation may contribute to the observed upregulation of Paraoxonase-1 (PON1), an esterase linked to high-density lipoprotein (Figure 5K). The fibroblast cell-derived secreted phosphoprotein-1 (SPP1), a protein associated with heart disease, was also found to be downregulated (Figure 5L).

The collagen-binding protein Von Willebrand factor (VWF) showed a significant downregulation (Figure 5M), following the downregulation of multiple ECM components. The integrin family, known for their collagen-binding properties, displayed varied responses, with ITGB2 and ITGAM being significantly downregulated, while ITGA11 showed a mild yet significant upregulation (Figure 5N-P). Additionally, the guanine nucleotide-binding subunit GNG7 was found to be up-regulated (Figure 5Q).

ECM components play a crucial role in immune regulation within the heart. Notably, the cathelicidin-like antimicrobial peptide (CAP11), Ig gamma-2 chain C region (p01862), and lactoferrin (LTF) were significantly downregulated in ARVC cohort (Figure 5S-U).

In the context of *pkp2* knock down, contain associated protein CNTNAP1 and CNTN1 were significantly upregulated after knockdown of *pkp2*. Neuronal growth factor 1 (NEGR1), significantly upregulated as well (Figure 5 V-X).

### Cardiomyopathy and Cardiac Muscle Contraction Proteins and metabolic remodeling in Guinea Pigs with PKP2 Knockdown

Proteomic analysis revealed differential regulation of several ECM proteins linked to cardiomyopathy and heart failure, including DPT, ITGA11, SPP1, P01862 are all recognized for their roles in the pathophysiology of heart failure. This led us to explore the status of other proteins integral to cardiac muscle contraction.

Chymase (CMA1), a serine protease predominantly located in cardiac mast cells and implicated in ECM remodeling of failing heart, was markedly upregulated in the right ventricular (RV) walls of PKP2 knockdown guinea pigs (Figure 6A). Concurrently, we noted an increase in Myosin Heavy Chain 6 (MYH6) and a decrease in Troponin 3 (TPM3) and Troponin 4 (TPM4) (Figure 6B-D), indicating a potential disruption in the structural integrity of cardiac muscle fibers.

**Figure 6.**
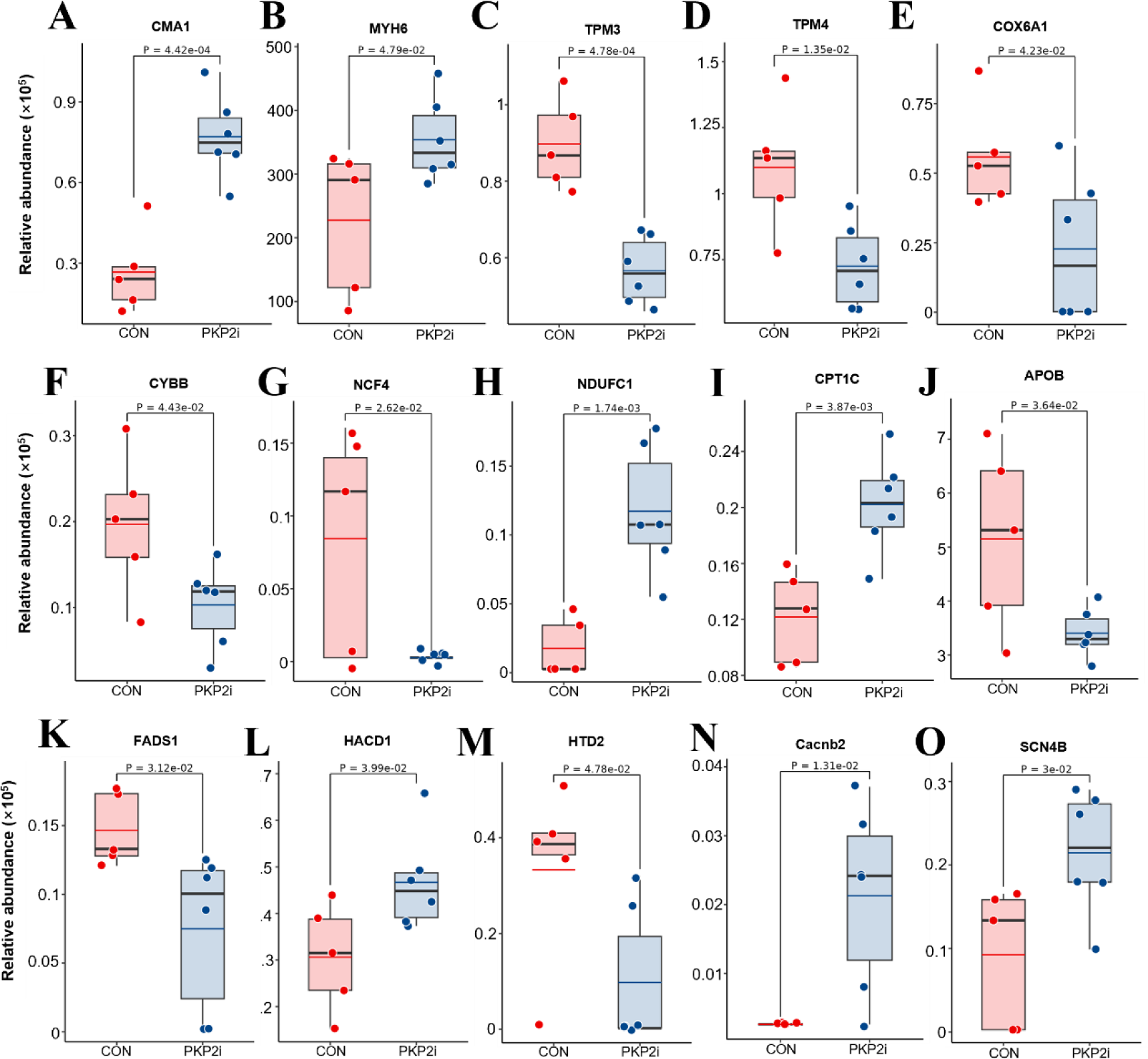
Altered cardiac muscle contraction and metabolic Protein Expression in PKP2-Deficient Right Ventricles. The differential expression analysis of cardiac muscle contraction and metabolic proteins in right ventricle samples from CON and ARVC animals, which are representative of PKP2 deficiency. Data are expressed as mean relative abundance ± standard deviation (SD), with n=5 for the CON group and n=6 for the PKP2i group. The Y-axis denotes the relative abundance of proteins, scaled by a factor of 10^5^. Statistical significance, as determined by P values, is indicated for each protein graph, highlighting the dysregulation of ECM components in the PKP2-deficient state.

These findings were complemented by a perturbation in proteins central to energy metabolism. Specifically, Cytochrome C oxidase subunit 6A1 (COX6A1) and cytochrome B-245 heavy chain (CYBB) were significantly downregulated (Figure 6E, F), which could impair mitochondrial respiration and ATP generation. In contrast, Neutrophil Cytosolic Factor 4 (NCF4), a component of NADPH oxidase, and NADH Dehydrogenase 1 Subunit 1 (NDUFC1) exhibited opposing regulatory patterns, with NCF4 downregulated and NDUFC1 upregulated (Figure 6G, H), suggesting a metabolic reconfiguration.

The observed alterations in energy metabolism were mirrored by changes in lipid metabolism. Carnitine Palmitoyltransferase 1C (CPT1C), which facilitates fatty acid transport into mitochondria, was upregulated (Figure 5I), while Apolipoprotein B (apoB) and Fatty Acids Desaturase 1 (FADS1) were downregulated (Figure 6J, K). Additionally, Very-Long-Chain (3R)-3-Hydroxyacyl-CoA Dehydratase 1 (HACD1) and Hydroxyacyl-Thioester Dehydratase Type 2 (HTD2) exhibited altered expression in the PKP2i group (Figure 6L, M), suggesting a shift in cardiac energy substrate utilization.

Building on the observed metabolic shifts, we also identified alterations in cardiac ion channels that could be consequential to the changes in energy metabolism. Notably, the Voltage-Dependent L-Type Calcium Channel Subunit Beta 2 (CACNB2) and Sodium Voltage-Gated Channel Beta Subunit 4 (SCN4B) were found to be significantly upregulated (Figure 6N, O).

### Metabolic Signatures and Pathway Alterations in PKP2 Knockdown Guinea Pigs Revealed by Plasma Metabolomics

To identify early diagnostic biomarkers for ARVC, we conducted a plasma metabolomics analysis using samples from CON and PKP2i guinea pigs.

Plasma samples from five animals in each group were analyzed using non-targeted metabolomics. Principal Component Analysis (PCA) distinguished a significant metabolic divergence between CON and PKP2i groups (Figure 7A). A total of 1,266 distinct metabolites were identified, with fatty acyls being the most prevalent (17.09%), followed by carboxylic acids and derivatives (11.97%), glycerophospholipids (11.11%), and benzene derivatives (8.55%) (Figure 7B). Differential metabolites were predominantly lipids and lipid-like molecules, constituting 33.05% of the total (Figure 7C). The dynamics of individual metabolite changes were visualized through volcano plots (Figure 7D) and circular heat maps (Figure 7E).

**Figure 7.**
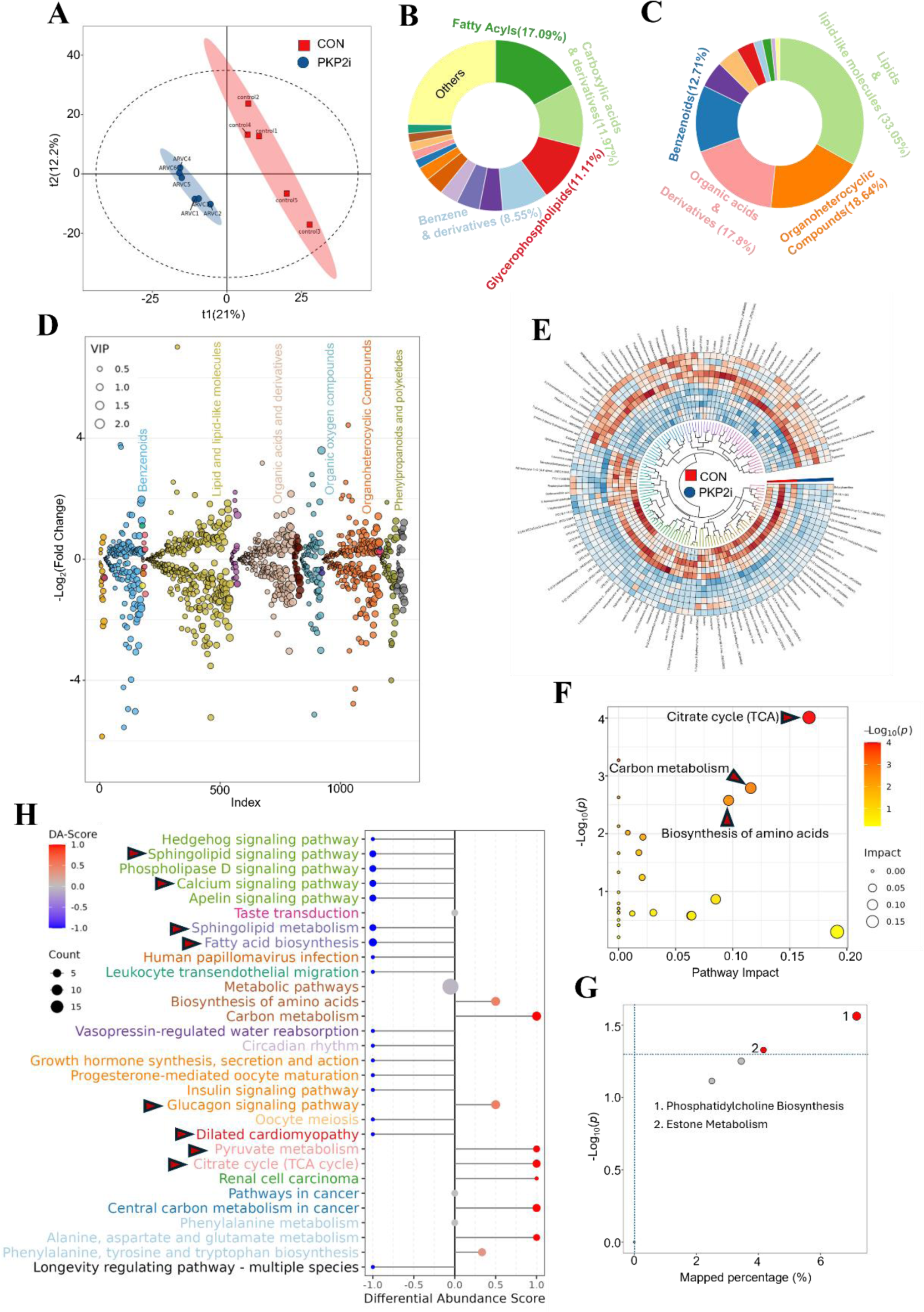
Metabolic Profile Alterations Induced by PKP2 Knockdown. (A) PCA plot comparing the metabolomic profiles of control (CON) and PKP2 knockdown (PKP2i) plasma samples, illustrating the metabolic divergence between the two groups. (B) Categorization of detected metabolites based on their chemical properties. (C) Classification of significantly differential metabolites, with a focus on the changes in chemical properties observed in PKP2i samples relative to CON samples. (D) Volcano plot depicting the differential metabolites, with the Y-axis representing the negative logarithm of the fold change (-log2 FC), allowing for the visualization of both the magnitude and significance of metabolic changes. (E) Circular heatmap illustrating the expression patterns of differential metabolites, with color intensity reflecting the level of expression and aiding in the identification of metabolic trends. (F) Pathway impact analysis using Betweenness centrality, which measures the influence of each pathway on the overall metabolic network. (G) The Small Molecule Pathway Database (SMPDB) primary pathway analysis, with bubbles sized proportionally to the significance (P<0.05) of the pathways affected by PKP2 deficiency. (H) KEGG pathway analysis, highlighting pathways of particular interest that are significantly impacted by PKP2 knockdown, as indicated by red arrows. Metabolomics data analysis was performed using Perseus, Excel, and R software, with enriched pathways considered significant at a p-value threshold of 0.05.

To elucidate the pathways influenced by these differential metabolites, we employed the Betweenness Centrality method to generate a pathway impact map. This analysis highlighted the Tricarboxylic Acid (TCA) cycle as the most significantly affected pathway in PKP2-deficient states (Figure 7F). Utilizing the Small Molecule Pathway Database (SMPDB) from the Human Metabolome Database (HMDB), we further analyzed the primary metabolic pathways. Phosphatidylcholine biosynthesis emerged as the most impacted pathway (Figure 7G). A Differential Abundance Score (DA-Score) was applied to rank and categorize the top 30 pathways, revealing upregulation of the TCA cycle, pyruvate metabolism, and glucagon signaling pathways, while fatty acid biosynthesis, sphingolipid metabolism and signaling pathways, calcium signaling pathways, and dilated cardiomyopathy pathways were downregulated (Figure 7H).

### Lipid Metabolome Reconfiguration and TCA Cycle Dysregulation in PKP2 Knockdown Guinea Pigs

Pathway analysis directed our attention to the significant changes in lipid and lipid-like molecules in the PKP2 knockdown (PKP2i) group. Specifically, oleic acid, palmitoleic acid, and isostearic acid were markedly downregulated (Figure 8A-C). Sphingomyelin species, including SM(18:0/16:1), and sphingosine 1-phosphate also showed a notable decrease (Figure 8D, E). In the context of phosphatidylcholines (PC), several species were dysregulated; PC(16:0/18:1), PC(17:0/18:1), and PC(18:0/20:3) were upregulated, while PC(18:1/14:0) and PC(18:0/20:3) were downregulated (Figure 8F-J). Conversely, multiple PC metabolites were significantly downregulated in the PKP2i group, such as phosphocholine, lysoPC(LPC)(18:2), LPC(16:0), LPC(18:1), and LPC(17:0) (Figure 8K-O). Despite the significant impact on PC, the levels of phosphatidylethanolamine (PE) remained relatively unchanged, with only PE(21:0/22:6) showing an upregulation (Figure 8P), indicating a shift in the PC/PE ratio.

**Figure 8.**
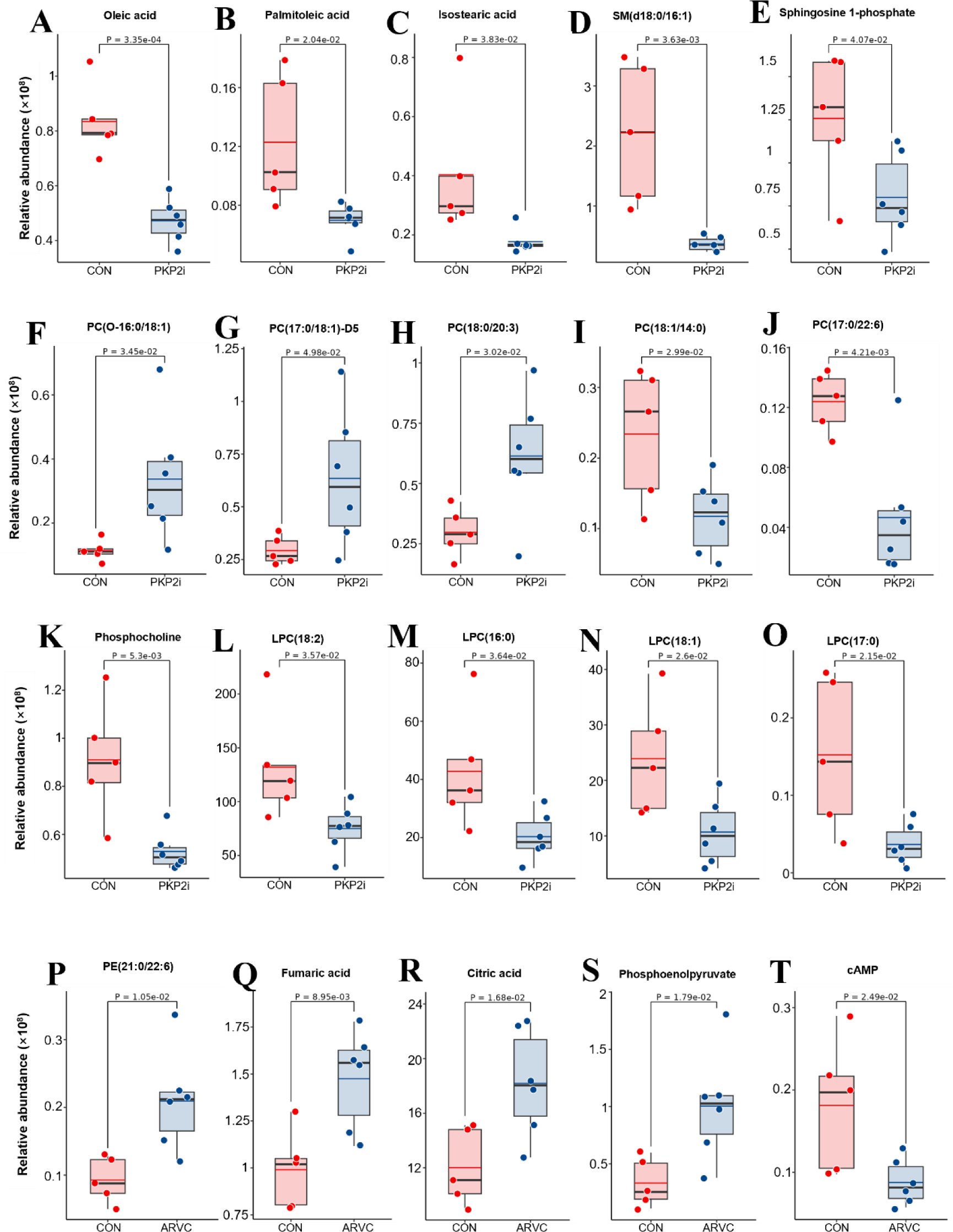
Changes in metabolites with PKP2 knockdown. The differential expression analysis of metabolites in plasma samples from CON and ARVC animals, which are representative of PKP2 deficiency. Data are expressed as mean relative abundance ± standard deviation (SD), with n=5 for the CON group and n=6 for the PKP2i group. The Y-axis denotes the relative abundance of proteins, scaled by a factor of 10^5^. Statistical significance, as determined by P values, is indicated for each protein graph.

Metabolites of the Tricarboxylic Acid (TCA) cycle, including fumaric acid, citric acid, and phosphoenolpyruvate, were found to be upregulated in the PKP2i group (Figure 8Q-S). Additionally, cyclic adenosine monophosphate (cAMP), a key molecule associated with dilated cardiomyopathy, was significantly downregulated in the PKP2-deficient state (Figure 8T).

## Discussion

ARVC is a significant contributor to SCD in youth^14^. However, the early pathological changes and progression of ARVC are not well understood^1^, partly due to the absence of animal models that fully replicate the human condition. To address this, we established an ARVC model in guinea pigs using AAV9-mediated delivery of PKP2 shRNA. This model exhibited several human-like symptoms, including increased cardiac lipid content, sudden death, and right ventricular (RV) enlargement, within four months^3^. The increased sensitivity observed in male guinea pigs compared to females parallels the male predisposition to ARVC, providing a valuable model for studying early ARVC pathogenesis^15^.

Our proteomic analysis of the RV myocardium has shed light on the molecular underpinnings of ARVC onset and the role of PKP2 in cardiac homeostasis. We identified a significant disruption in ECM homeostasis following PKP2 knockdown, potentially due to the upregulation of Foxo3, a key transcription factor known to promote ECM degradation. The activation of Foxo3 has been reported to enhance the degradation of ECM components, such as fibronectin (FN1), in endothelial cells and cardiac fibroblasts^16^, vascular smooth muscle^17^ and cardiac fibroblast^18^. In our study, the upregulated Foxo3 was associated with a significant downregulation of FN1 and fibrinogens, indicating a potential mechanism for ECM remodeling in ARVC.

Similar with FN1, another ECM component, ECM1 also negatively related with Foxo3 in a disease named Lichen Sclerosus^19^, same with our observation in this ARVC model. Additionally, we observed downregulation in ECM binding proteins, including VWF and integrin family proteins, with the exception of ITGA11, which showed a mild but significant increase in the PKP2 knockdown group. These changes suggest a broader impact on the ECM structure and its associated proteins, which could have implications for cardiac function.

### EMC remodeling or disorder also affected some functional proteins

The protease inhibitors A2M and several serpin family proteins were found to be significantly decreased, indicating a disorder in the ECM region’s protein composition. Immunity-related ECM proteins such as CAP11, P01862, and LTF were also significantly downregulated. CAP11, an antimicrobial peptide, and P01862, an immune globin gamma-2 chain C region, are known to play roles in immune defense. LTF, an iron transport protein with antibacterial activity, has been associated with lipid levels in blood^20^, suggesting a potential regulatory role in both cardiac and systemic lipid metabolism.

The PI3K-Akt pathway, which is known to regulate cardiac metabolism^21^, may also be implicated in the metabolic alterations observed in our study. Several differentially expressed ECM proteins, including ITGA11, FN1, SPP1, VWF, P01862, and GNG7, are involved in the regulation of the PI3K-Akt pathway. The downregulation of these proteins in the PKP2 knockdown group could explain the upregulation of Foxo3, as it is a primary target of Akt-mediated inactivation^22,23^. Dysregulation of the PI3K-Akt signaling pathway could contribute to the metabolic changes observed in PKP2-deficient RVs.

The upregulation of cardiac CPT1C when PKP2 deficient suggests enhanced fatty acid oxidation and a shift in energy substrate utilization in the heart. Changes in other lipid metabolic proteins, such as FADS1, HACD1, and HTD2, further support this shift in energy metabolism.

Energy substrate shifts are known to be a significant cause of heart failure^24–26^. Therefore, we hypothesize that alterations in metabolic proteins contribute to ARVC-related heart failure. Proteins involved in cardiac muscle contraction were also found to be disordered in the PKP2 knockdown state, with increased MYH6 and decreased tropomyosin 3 and 4, indicating a disrupted architecture of cardiac muscle fibers.

Additionally, the link between ECM and mitochondria is already well established in skeletal muscle and C. elegans^27,28^. This raises the possibility that PKP2 deficiency may also disrupt mitochondrial-ECM crosstalk in the heart, contributing to the mitochondrial morphological anomalies we observed.

Notably, key mitochondrial respiratory chain proteins, COX6A1 and CYBB, which facilitate electron transfer in complexes IV and III respectively^29,30^, were found to be downregulated. This downregulation likely indicates a compromised ATP-generating capacity.

Conversely, the upregulation of NDUFC1, a subunit of complex I, was observed in the PKP2 knockdown group. Given that complex I is involved in NAD+ production, an essential factor for fatty acid oxidation, this upregulation may be a compensatory response to the increased fatty acid oxidation signaled by the upregulation of CPT1C.

Plasma metabolomic analysis revealed a decrease in fatty acids alongside an accumulation of fumaric acid, citric acid, and phosphoenolpyruvate, suggesting an upregulation of fatty acid oxidation coupled with a suppression of oxidative phosphorylation. These findings point to a metabolic shift that could have significant implications for cardiac energy metabolism and function in the context of PKP2 deficiency. Furthermore, our plasma metabolomic analysis revealed significant alterations in the levels of several PC species, with a notable downregulation of LysoPCs and phosphocholine. In contrast, among the PE species, only PE(21:0/22:6) demonstrated a change in the context of PKP2 deficiency. This observation leads us to hypothesize that the PC/PE ratio may be altered in this model. The perturbation of the PC/PE ratio has been implicated as a biomarker in various metabolic disorders, including chronic organ failure^31^. Consequently, future research should investigate the potential association between alterations in the plasma PC/PE ratio and the development of chronic failure related to ARVC.

Further, certain individual proteins identified in our study suggest potential etiological links to ARVC. DPT, which is known to be upregulated in myocardial infarction (MI) zones, and SPP1, a protein secreted by cardiac fibroblasts and implicated in heart failure across various cardiomyopathies, exhibited contrasting expression patterns in the context of PKP2 deficiency. Specifically, DPT was upregulated, while SPP1 was downregulated in the PKP2-deficient state. These findings warrant further investigation into the role of these proteins in the pathogenesis of ARVC and their potential as therapeutic targets.

In summary, the guinea pig model of ARVC, characterized by the absence of overt heart failure and normal ECG findings, captures an early disease stage, offering a unique window into the initial pathogenic processes of ARVC. This model has revealed that PKP2 deficiency leads to RV ECM dysregulation, metabolic remodeling, and respiratory chain alterations, which may precede the clinical manifestations of heart failure associated with ARVC. Unraveling the intricate mechanisms that link PKP2 suppression to ECM imbalance and energy metabolism disruptions will be a critical area of future research. Despite the normal ECG results, the occurrence of sudden death in the model underscores the potential for malignant arrhythmias to remain undetected, highlighting the need for a more nuanced understanding of the interplay between ECM remodeling, metabolic disturbances, and electrophysiological consequences of PKP2 knockdown.

This study provides a comprehensive overview of the molecular and metabolic changes in the RV, as well as the pathways affected by PKP2 deficiency, thereby shedding light on the early molecular underpinnings of ARVC. However, further functional studies are essential to confirm the roles of the identified proteins and pathways in the disease process.

While the guinea pig model offers certain advantages over murine models, it is important to recognize that it may not fully mimic the complex pathology of human ARVC. The physiological differences between guinea pig and human hearts must be considered when interpreting the findings and translating them to clinical scenarios. Despite these limitations, this work contributes significantly to the understanding of early ARVC pathogenesis and sets the stage for future investigations into potential therapeutic targets.

## Methods and Material

### Animal Models

Sixteen male and sixteen female guinea pigs (250g) were purchased from Shuangxin Experimental Animals Co. Ltd, China. All animals were maintained in the Large Animal Facility at the Yunnan Agricultural University in a 12-hour light/dark cycle at RT 23-26 ℃. After one week of acclimatization, animals were randomly assigned to receive intravenous injection of 1 x 10^9^ AAV9 viral particles carrying either PKP2 shRNA (n=16) or non-targeting control shRNA (n=16). After 2 weeks of injection, animals were subject to electrocardiogram (ECG) and Echocardiographic. 4 months later, guinea pigs were sacrificed, and the left and right ventricle were sampled.

### AAV Vector Construction

AAV9 vectors expressing guinea pig PKP2 shRNA or non-targeting control shRNA were constructed and packaged into viral particles by OBiO Technology Corp., Ltd (Shanghai, China). The PKP2 shRNA target sequence GGATGTACTTGTCCTTGAT was selected after testing efficacy of three candidates. The non-targeting control sequence was CCTAAGGTTAAGTCGCCCTCG.

### Tissue Sectioning

After sacrificing, hearts were isolated, separated into left and right ventricles, and fixed in formalin at 4 ℃ for 48 hours. Then samples were Dehydrate the tissues in EtOH baths. Next, the tissues were cleared in xylene. Then tissues were embedded into paraffin, sectioned at 6 μm.

### HE and staining

Paraffin-embedded tissue sections were deparaffinized and rehydrated to distilled water. Slides were stained with Mayer’s hematoxylin solution for 1 minute to label nuclei, rinsed in tap water, and immersed in PBS for blueing. After washing in distilled water, slides were counterstained with alcoholic eosin Y solution for 1 minute. Finally, sections were dehydrated through increasing concentrations of ethanol and cleared in xylene prior to mounting.

### Sirius Red Staining

Following deparaffinization and hematoxylin nuclear staining, sections were incubated with 0.1% Picrosirius Red solution (Sigma-Aldrich) for 1 hour at room temperature. Slides were then rinsed twice with 0.5% acetic acid, dehydrated in increasing ethanol concentrations, cleared in xylene, and cover slipped.

### Transmission Electron Microscopy

Right ventricular tissues were fixed in TEM buffer (Servicebio) for 2 hours followed by post fixation in 1% osmium tetroxide. Samples were then dehydrated through a graded ethanol series and embedded in EMbed 812 resins (Electron Microscopy Sciences). Ultrathin sections of 60 nm thickness were cut using a Leica UC7 ultramicrotome and collected on 150 mesh copper grids. Prior to visualization, grids were stained sequentially with uranyl acetate and lead citrate solutions to enhance contrast. Sections were imaged using a Hitachi HT7800 transmission electron microscope.

### Electrocardiogram (ECG)

Guinea pigs were anesthetized by intraperitoneal injection with 0.5 mg/kg of Pentobarbital sodium. Surface six-lead ECG was recorded using standard limb lead placements. Tracings were acquired for ≥5 minutes after stabilization.

### Echocardiographic Evaluation

Transthoracic echocardiography was performed monthly under anesthesia to evaluate cardiac structure and function. Images were acquired in long-axis, short-axis, and apical four-chamber views using a Vivid E95 system (GE Healthcare) equipped with an 8-18 MHz transducer. M-mode, 2D, and pulsed-wave Doppler modalities were utilized. Quantitative structural parameters included left and right atrial diameters, right ventricular outflow tract dimensions, right ventricular basal and median diameters, and left ventricular wall thicknesses. Functional parameters consisted of right ventricular fractional area change, tricuspid annular plane systolic excursion, myocardial performance index, left ventricular ejection fraction, and fractional shortening. Echocardiographic measurements were obtained according to standardized protocols^32^.

### Sampling and proteomics

Right ventricular tissues were flash frozen in liquid nitrogen immediately after isolation. DIA (Data independent acquisition) quantitative proteomics was performed. Proteins samples were enzymatically digested into peptides and chromatographically separated using a *Vanquish Neo UHPLC system* (Thermo Scientific). Then the separated peptides were performed DIA mass spectrometry analysis using an Orbitrap *Astral* mass spectrometer (Thermo Scientific). Results were analyzed by MSFragger 3.4 software using the UniProt Cavia porcellus database: uniprot-Cavia porcellus (Guinea pig)-25662-20221222.fasta (https://www.uniprot.org/uniprotkb?query=(taxonomy_id:10141)).

### Metabolomics assay

Samples were weighed before the extraction of metabolites and dried lyophilized were ground in a Grinding Mill. Metabolites were extracted using 1 mL precooled mixtures of methanol, acetonitrile and water (v/v/v, 2:2:1).

Metabolomics profiling was analyzed using a UPLC-ESI-Q-Orbitrap-MS system (UHPLC, Shimadzu Nexera X2 LC-30AD, Shimadzu, Japan) coupled with Q-Exactive Plus (Thermo Scientific, San Jose, USA). The raw MS data were processed using MS-DIAL for peak alignment, retention time correction and peak area extraction. The metabolites were identified by accuracy mass (mass tolerance < 10 ppm) and MS/MS data (mass tolerance < 0.02Da) which were matched with HMDB, massbank and other public databases and our self-built metabolite standard library. In the extracted-ion features, only the variables having more than 50% of the nonzero measurement values in at least one group were kept.

### Triglyceride (TAG) assay

Triglyceride (TAG) levels in right ventricular (RV) tissue were measured using a commercial assay kit (Applygen Technologies Inc.). Briefly, 50 mg of RV tissue was weighed, minced, and lysed in 1 ml of lysis buffer by vortexing. The homogenate was centrifuged at 2000 rpm for 5 min and the supernatant collected. Supernatant samples were mixed with the kit-provided R1 and R2 buffers and TAG concentration measured by absorbance at 570 nm using a microplate reader, according to the manufacturer’s protocol.

### Western Blot

Anti-guinea pig PKP2 antibody was produced by Abclone Ltd. Right ventricular lysates were prepared using RIPA buffer (Santa Cruz Biotechnology). 30 μg protein was separated by western blot. Blots were probed for PKP2 (1:1000) and GAPDH (Abclone, 1:2000). Signals were quantified by densitometry using an Imaging System (Bio Technology, China).

### Statistical Analysis

Echocardiographic Evaluation was conducted 8 animals of each group. Proteomics experiments were conducted in triplicate. Data are expressed as the mean ± SEM. Statistical differences were analyzed using GraphPad Prism software (GraphPad Software Inc., San Diego, CA). Statistical significance was then determined by unpaired two-tailed Student’s t-test with a threshold of significance set at p<0.05.

## Acknowledgements

We thank OBiO Technology (Shanghai) Corp., Ltd for AAV9 and shRNA package construction. We thank Shanghai Bioprofile Technology Company Ltd for proteomics study and bioinformation analysis. This work was supported by Yunnan Fundamental Research Projects (Grant NO. 20221AS070081), National Natural Science Foundation of China (Grant NO. 32060206) and veterinary Public Health Innovation Team of Yunnan Province (Grant NO. 202105AE160014).

## Author contributions

Haizhen Wang and Fei Sun performed study concept and design; Rui Song and Haiyan Wu performed Echocardiographic evaluation of right ventricular structure and sampling for proteomics analysis. Jingning Yu and Wenhui Yang performed the ECG analysis, western blot analysis and sampling. Wenjun Wu raised all the guinea pigs in this study. Haizhen Wang and Fei Sun revised the manuscript. All authors read and approved the final paper.

## Ethics approval and consent to participate

All aspects of this study were approved by the Experimental Ethics Committee of Kunming Medical University.

