## supplemental figures for "Multiomics Analysis Reveals Extensive Remodeling of the Extracellular Matrix and Cellular Metabolism Due to Plakophilin-2 Knockdown in Guinea Pigs"

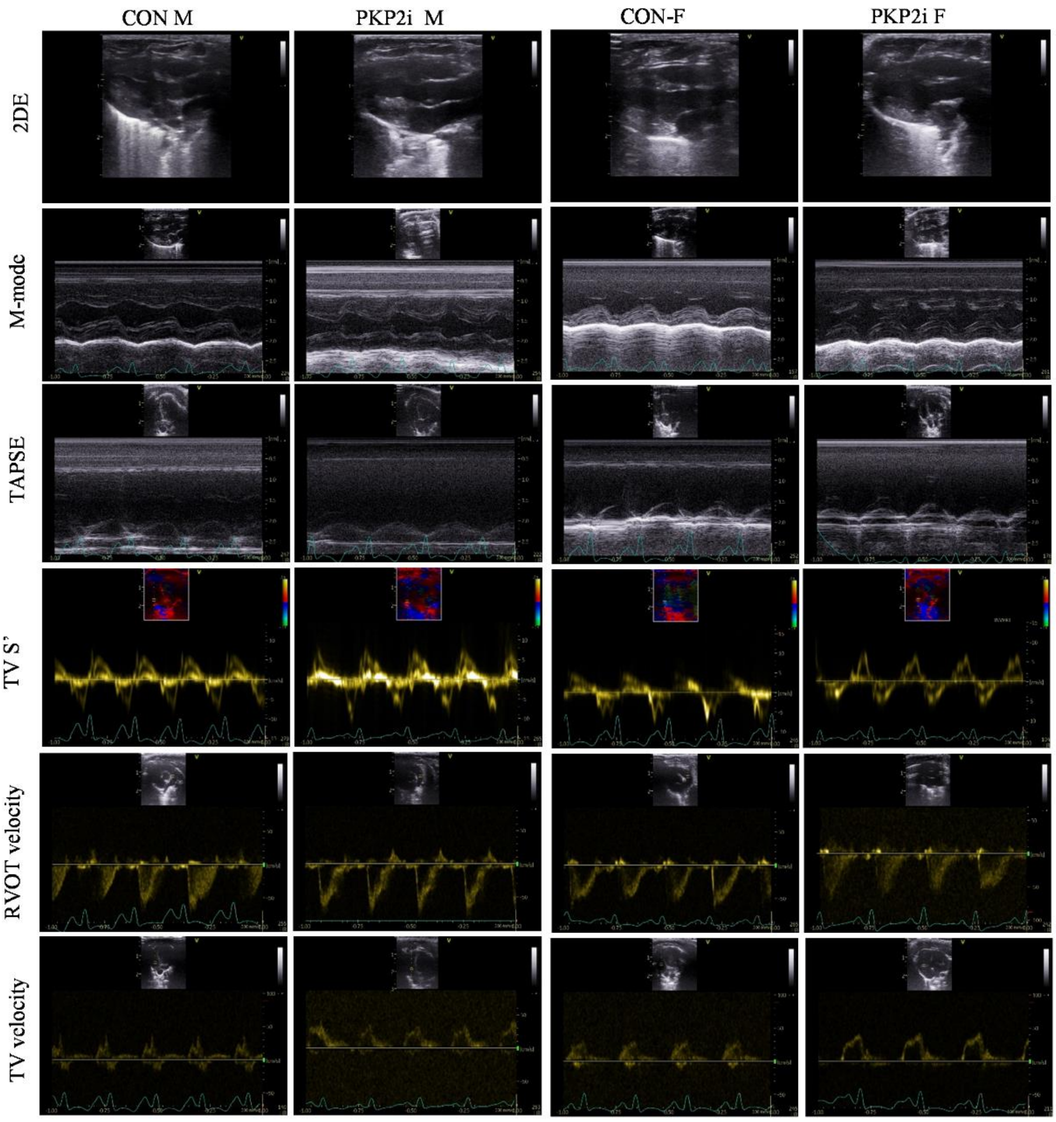


Supplementary Figure 1. Left ventricle function measurement by Echocardiography.

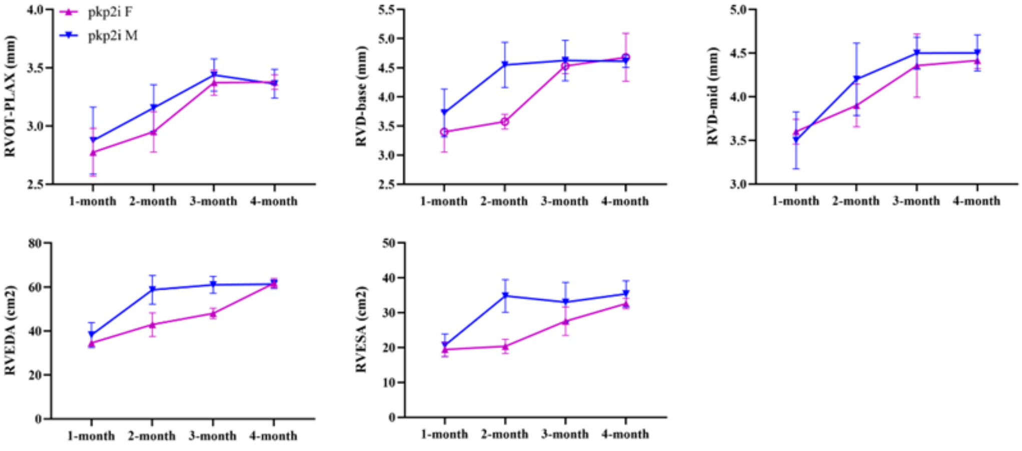


Supplementary Figure 2. Compare of male and female of RVOT-PLAX, RVD-base,

RVD-mid, RVEDA and RVESA.
